# Distinct patterns of connectivity with motor cortex reflect component processes of sensorimotor learning

**DOI:** 10.1101/2023.07.01.547344

**Authors:** Corson N. Areshenkoff, Anouk J. de Brouwer, Daniel J. Gale, Joseph Y. Nashed, J. Randall Flanagan, Jonathan Smallwood, Jason P. Gallivan

## Abstract

Sensorimotor learning is supported by multiple competing processes that operate concurrently, making it a challenge to elucidate their neural underpinnings. Here, using human functional MRI, we identify three distinct axes of connectivity between the motor cortex and other brain regions during sensorimotor adaptation. These three axes uniquely correspond to subjects’ degree of implicit learning, performance errors and explicit strategy use, and involve different brain networks situated at increasing levels of the cortical hierarchy. We test the generalizability of these neural axes to a separate form of motor learning known to rely mainly on explicit processes, and show that it is only the Explicit neural axis, composed of higher-order areas in transmodal cortex, that predicts learning in this task. Together, our study uncovers multiple distinct patterns of functional connectivity with motor cortex during sensorimotor adaptation, the component processes that these patterns support, and how they generalize to other forms of motor learning.

---

Many of our daily actions stem from the interplay of different mental processes that operate concurrently to achieve our intended goals. Broadly, these mental processes can be distinguished by the nature of their cognitive demands: Explicit processes involve conscious mental operations, often requiring our directed attention and deliberate thought; by contrast, implicit processes operate autonomously, operating beneath the threshold of our conscious awareness [1]. Extensively examined within the realms of psychology and neuroscience, these processes often operate in tandem to accomplish a wide range of tasks, from learning a foreign language to playing a musical instrument or driving a car [2, 3, 4].

In recent years, there has been significant interest in understanding how these two separate processes support motor learning. For decades, motor learning was widely believed to constitute a singular, implicit process of the sensorimotor system [5, 6]. However, recent behavioral and computational work has demonstrated that motor learning is actually supported by at least two separate learning systems that operate in parallel: a slow-acting implicit system that adapts gradually, and a fast-acting explicit system that adapts rapidly [7, 8, 9, 10, 11, 12, 13], and that may depend on brain areas located outside the sensorimotor system. Notably, as performance during learning can reflect the summed contribution of both processes, the ability to characterize their underlying neural systems has remained a challenge. This is because these processes not only need to be disentangled from each other, but also because they must be dissociated from neural activity related to other facets of performance (e.g., sensory feedback related to errors).

Previous research has often framed implicit sensorimotor learning within optimal feedback control theory, with general agreement that such learning reflects the updating of an internal model that links motor commands to sensory outcomes. This form of learning is thought to be supported by a network of cerebellar and sensorimotor cortical regions, crucial for the sensory-guided control of movement [14, 15, 16]. In contrast, the neural foundations of explicit learning are considerably less well understood, and may well depend upon regions of cortex that serve more general functions related to cognition, such as association cortex [17]. For example, explicit processes such as strategy use have been associated with the activity of prefrontal cortex [18]; however, it is unclear whether this activity reflects the *use* of explicit strategic processes *per se* or the attendant outcomes of using those processes (e.g., the resulting change in performance errors). This is an important neural distinction for understanding how we guide complex behavior, since explicit processes are often brought to bear at critical moments during learning, and can result in a rapid reduction in visuomotor errors [13].

Here, using functional MRI to study whole-brain patterns of functional connectivity with motor cortex, we sought to disentangle these different processes associated with sensorimotor learning. We used subjects’ learning behavior in combination with their reports of explicit strategy use to identify the relative contributions of explicit and implicit processes to learning, as well as the trajectory of errors that they make during the task. These behavioral measures were used to identify three distinct axes of whole-brain connectivity with the motor cortex that emerged during early learning and that corresponded to subjects’ implicit learning, their performance on the task (error rate), and their use of explicit strategic processes, respectively. We found that these separable neural axes were approximately situated at different hierarchical levels of cortical organization; implicit learning was associated with the connectivity of areas in superior parietal and premotor cortex, whereas subjects’ visuomotor performance was associated with the connectivity of a subset of regions in the frontoparietal control network. Notably, we found that explicit learning was associated with the connectivity of a subset of areas positioned at the apex of the cortical processing hierarchy, in the default mode network. We tested the generalizability of these separate neural axes by studying sensorimotor learning in a different task – performed in the same subjects – in which abrupt changes in performance are known to rely mainly on explicit strategic processes [19, 20]. Together, our findings dissociate three distinct patterns of connectivity with motor cortex that correspond to separate neural processes that operate concurrently during sensorimotor learning.

## 1 Results

### 1.1 Learning behaviour during sensorimotor adaptation

To study the neural processes that support sensorimotor learning, we had subjects (N=36) perform a visuomotor rotation task [VMR; 21] in the MRI scanner. In this task, subjects were required to move a cursor, representing their right finger position, to intercept a target that could appear in one of eight locations on a circular ring, using an MRI-compatible touchpad (Figure 1. After a baseline block (40 trials), subjects performed a learning block (160 trials) in which the correspondence between finger motion on the touchpad and movement of the cursor was rotated clockwise by 45*^◦^*. Thus, subjects needed to learn to adapt their movements by applying a 45*^◦^* counter-clockwise rotation in order to hit the target. At the conclusion of each trial, subjects received visual (error) feedback consisting of both the target location, as well as the position of their cursor on the target ring. Following the learning block, subjects performed 16 “report” trials [13], in which they were asked to indicate their aim direction using a dial, controlled by their left hand, prior to executing a target-directed movement via their right hand. These report trials were presented at the end of the task, after learning had taken place, in order to avoid drawing attention to the nature of the visuomotor perturbation and thus biasing subjects’ learning behaviour [22]. Critically, these report trials provided us with an direct index of subjects’ explicit knowledge of, and strategy in counteracting, the visuomotor rotation [13].

**Figure 1:**
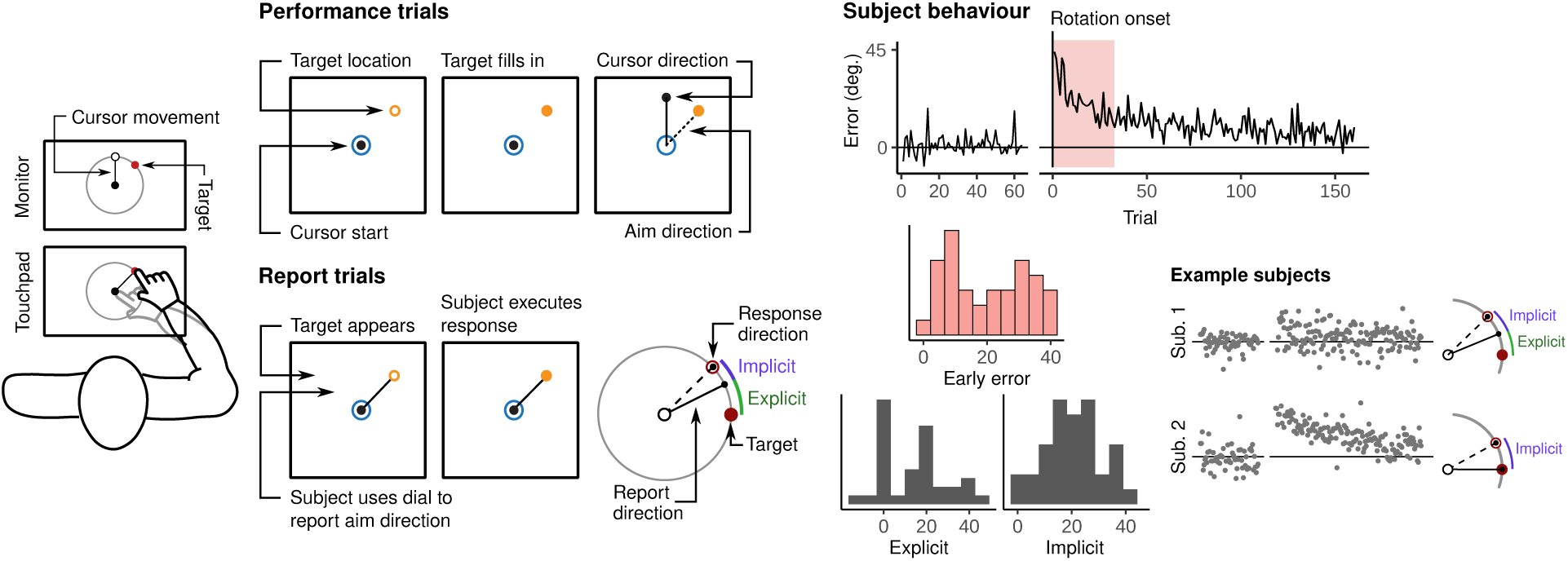
Visuomotor rotation task and subject behaviour – (left) Subjects were cued to move a cursor to a target appearing on a circular array using an MRI compatible touchpad. During learning trials, the cursor moved at a 45*^◦^* angle relative to the movement on the touchpad, so that subjects’ had to learn to adjust their aim direction in order to contact the target successfully. After learning, subjects performed a set of report trials in which they used a dial to indicate their aim direction prior to executing the movement. Using these report trials, we derived estimates of subjects’ total explicit and implicit learning during the task. (**Right**) Our analysis focused on functional connectivity during the early learning period, when the reduction in error is most rapid and explicit processes are believed to be most prominent. Histograms at bottom left show the distribution of subject errors during the early learning period (in red), and explicit and implicit learning components across subjects (in dark grey). Two example subjects are shown at right.

Behavioral data associated with the VMR task are shown in (Figure 1. We found that subjects, on average, were able to successfully learn the VMR task by aiming their hand in a direction that counteracted the visuomotor rotation (Fig 1, left). However, it is important to note that these results obscure significant inter-subject differences in the rate at which subjects’ were able to reduce their errors. For instance, Fig 1 (right) highlights the learning curves for two example subjects: One individual who was able to reduce their visuomotor errors fairly rapidly during early learning (top plot, Subject 1), and another individual who was able to only gradually reduce their errors during this same period (bottom plot, Subject 2). Notably, as indicated by the declarative report trials performed at the end of learning (see rightmost graphics), the more rapid learner expressed some degree of explicit knowledge about the rotation (i.e., aimed counter-clockwise of the target) whereas the more gradual learner did not (i.e., aimed directly at the target). On this point, we observed a large degree of variability in subjects’ aiming directions during these report trials, ranging from zero explicit knowledge of the rotation (wherein subjects reported aiming directly at the target, or 0*^◦^* explicit report) to complete compensation (i.e., wherein subjects aimed 45*^◦^* to fully counteract the imposed rotation; see histogram). Although the rapid reduction in errors during early learning are believed to reflect an explicit re-aiming process [13, 22], performance *per se* during this period reflects the summed contribution of both explicit and implicit processes, and so early learning cannot be taken as a direct proxy for either process.

To identify patterns of neural activity specific to subjects’ degree of implicit and explicit learning, as well task performance, we derived well-established measures of these different learning components for each participant [13, 22]. Consistent with prior work [13], we defined explicit learning as the mean difference between subjects’ reported aim direction and the target location, and defined implicit learning as the mean difference between subjects’ reported and actual aim directions. Finally, we defined task performance as subjects’ average error during the early learning period (defined as the first 100 imaging volumes, or 34 trials, of the task). Even though we did not identify any relationship between subjects’ use of explicit strategies (from report trials) and early learning performance (*r* = *−*0.15, *t*_34_ = *−*0.91, *p* = .37), we nevertheless included subjects’ early error scores as a regressor in both our analyses of subjects’ explicit and implicit learning data, thus controlling for any putative performance-related effects (see Methods for further details on this approach).

### 1.2 Connectivity-based approach for studying sensorimotor adaptation

To examine the neural bases of the different component processes identified in the previous section, we examined learning-dependent changes in functional connectivity between the primary motor cortex and the rest of cortex, striatum, and cerebellum (Figure 2, left). By constraining our analysis to bipartite functional interactions with primary motor cortex, we were specifically interested in assessing how various regions in the cortex, striatum, and cerebellum, modulate their connectivity with cortical areas positioned in the final motor output pathway that produce motor commands resulting in the measured behavioral adjustments. For each brain region, we extracted fMRI timecourse activity from different atlas parcellations of the cerebellum, striatum and all of cortex (see Methods), and studied functional connectivity between these brain regions with the average activity of a subset of 4 regions in the left (contralateral) hemisphere comprising the hand motor region [belonging to the left Somatomotor A network; 23]. For each subject, we estimated covariance networks during the baseline, early learning and late learning periods (each defined as a epoch of 100 imaging volumes), allowing us to describe changes in connectivity over the course of the task. Given that a large body of recent literature suggests that functional connectivity is dominated by static, subject-level differences that can obscure any task-related variance [26], we first centered subjects’ covariance matrices to align their mean covariance across resting state, baseline, and learning [see Methods for details, and 27, for a more thorough description].

**Figure 2:**
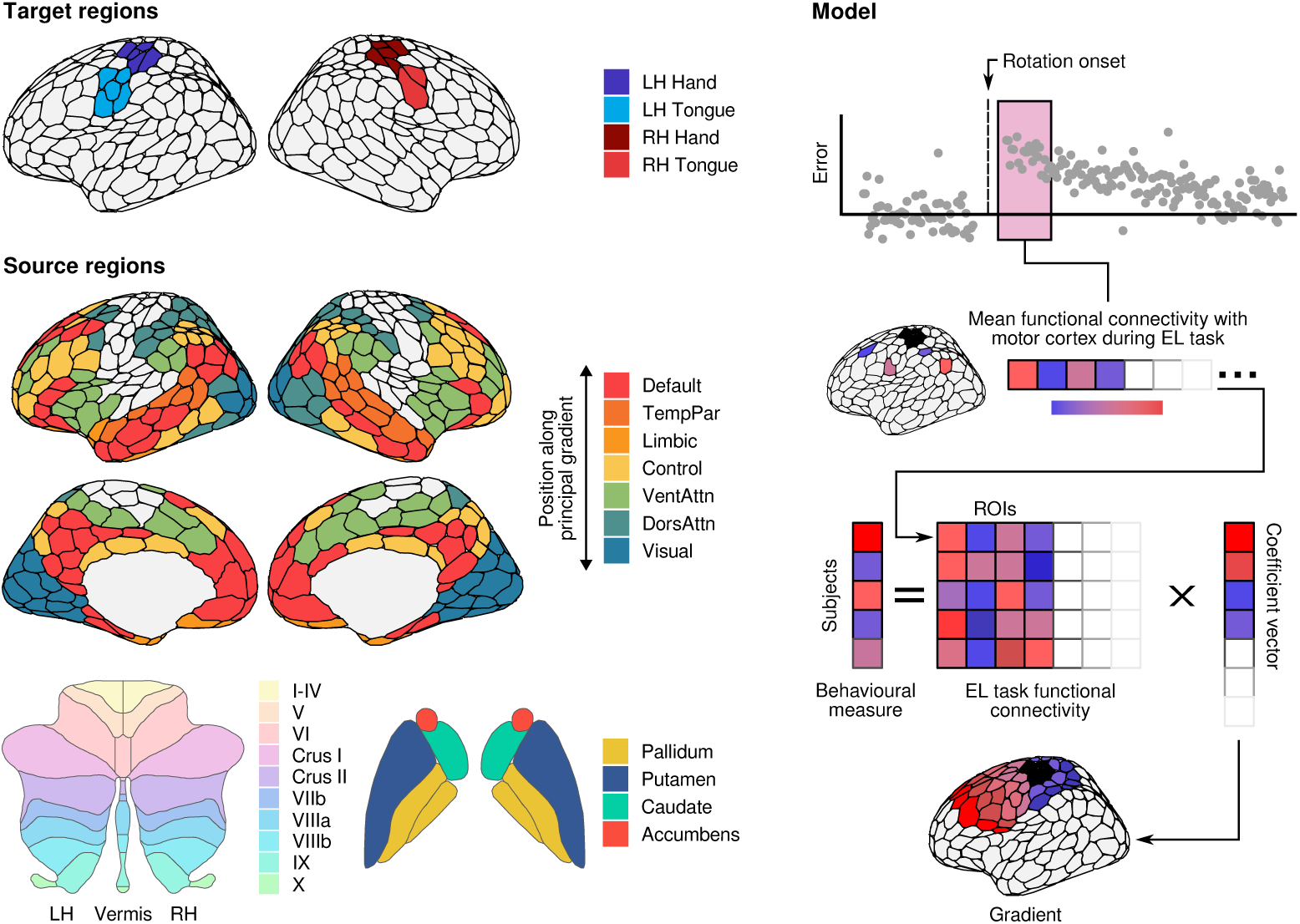
Brain network identification and predictive model – (left) We studied functional connectivity between primary motor cortex and the rest of cortex, striatum, and cerebellum. Cortical networks were derived using the 400 region parcellation of [23] and given network assignments by [24]. Note that, while we use the 17 network solution for the majority of our analyses, some of our results are reported at a coarser level (e.g. “Default mode” for Default A, B, and C). These coarse labels are, in general, nonequivalent to the 7 network solution. Cerebellar regions were identified using the atlas of [25]), while striatal regions were extracted from the Harvard-Oxford atlas. Motor cortex parcels were identified as belonging to either hand or tongue areas of the left and right hemispheres based on the reported findings of [23]. (**Right**) We used a set of penalized regression models (see Methods) to predict subjects’ explicit learning, task performance, and implicit learning from patterns of whole-brain functional connectivity with primary motor cortex. The coefficients from each of the fitted models produces a set of brain maps which we hereby refer to as the *explicit*, *performance*, and *implicit* neural axes.

Using the patterns of functional connectivity across regions extracted from the early learning period, we used a group-penalized ridge regression model to predict subjects’ explicit and implicit learning, as well as their performance errors [28, ; see Figure 2 right, and see methods]. We initially focused our analysis on this early learning period as this is when the contributions of explicit processes to learning are thought to be most prominent [13]. We used the region-wise coefficients from this model to create whole-brain connectivity maps reflecting covariance with motor cortex during early learning, which we hereby refer to as the Explicit, Implicit and Performance neural axes (Figure 3). Note that we inverted the sign of subjects’ error in the derivation of the performance axis, so that greater loadings were associated with lower error (i.e. better performance). For each axis, network specific penalties were tuned by leave-one-out cross-validation (cross-validated *R*^2^: *R*^2^ = 0.29, *R*^2^ = 0.82, *R*^2^ = 0.64).

**Figure 3:**
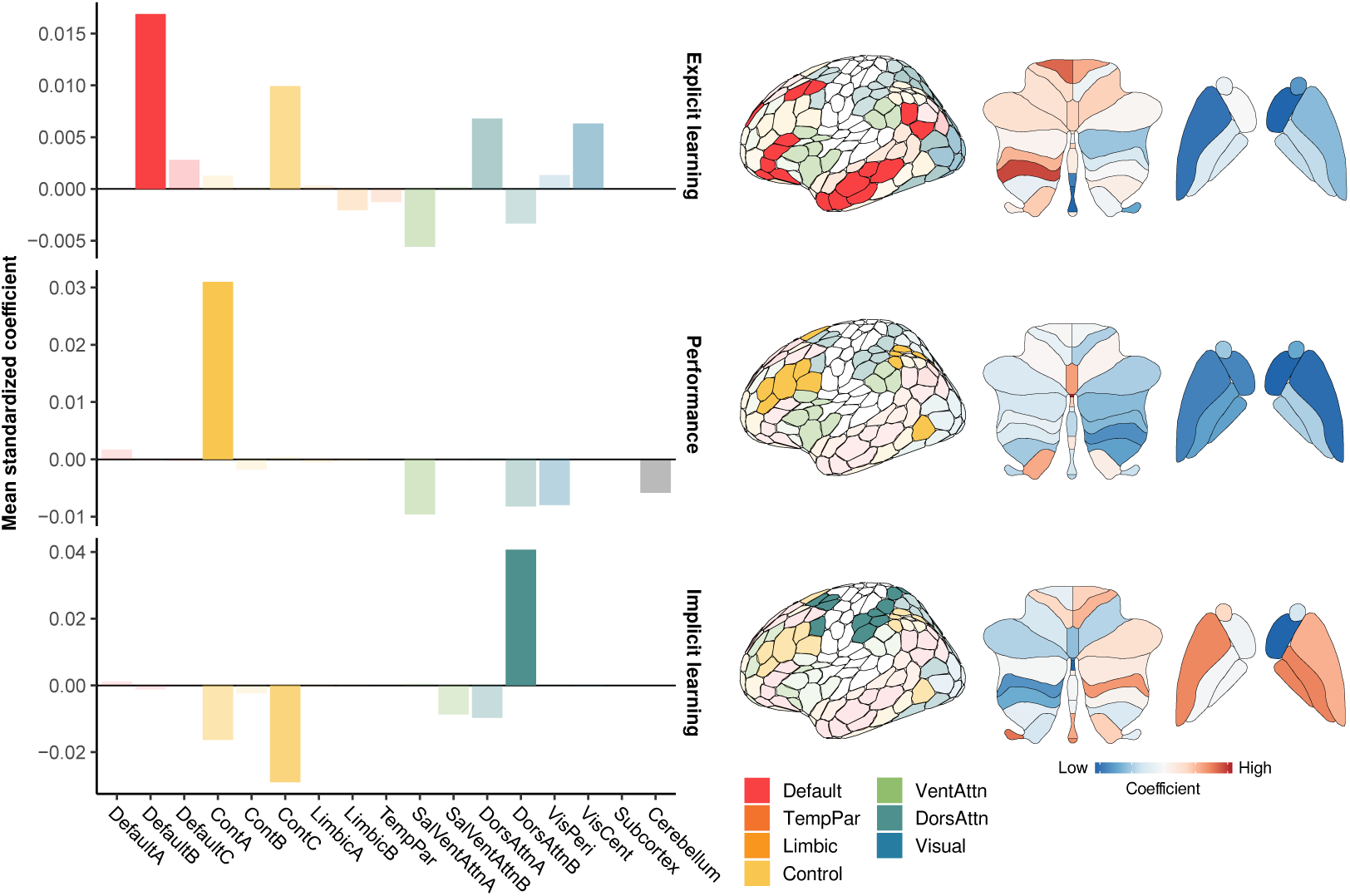
Explicit, performance, and implicit neural axes –. Barplots show mean coefficients from our penalized ridge regression model for each subnetwork of the 17 network assignments of [24]. The most prominent subnetworks for each neural axis are displayed on the adjacent cortical surfaces. Loadings for cerebellar and striatal regions are shown at the right; note that colormaps for each surface are scaled individually.

### 1.3 Separate neural axes are associated with different component processes of adaptation

Within the Implicit neural axis, we observed the highest regional loadings in the dorsal attention network (DAN), and specifically the DAN-B subnetwork, comprising superior parietal and premotor regions known to support the sensory-guided control of movement [14, 15, 16]. By contrast, within the Performance neural axis, we observed the highest regional loadings within the Control network, and specifically the Control-A subnetwork, mainly comprising the dorsolateral prefrontal cortex (DLPFC), inferior parietal lobe (IPL) and anterior cingulate cortex (see barplots in Figure 3). In the sensorimotor adaptation literature, the lateral prefrontal cortex has been shown to increase its activity during early learning [29, 30, 18], and thus it has been suggested to play a key role in implementing explicit strategies during learning [31, 32]. However, on this point it noteworthy that, within the Explicit neural axis, we actually observed the highest regional loadings within the default mode network (DMN); specifically, within the DMN-B subnetwork, comprising higher-order regions in the middle and inferior frontal gyrus, angular gyrus and anterior and middle temporal cortex (see barplots in Figure 3). In the sensorimotor adaptation literature, these DMN areas have not been commonly identified as having an important role during motor learning, though this likely reflects the fact that prior studies have not sought to directly isolate the use of explicit processes (e.g., through verbal reporting) during fMRI testing.

Note that, due to the sparsity of the coefficients returned by our models, we found that most cortical networks loaded only on a single axis (e.g., the DAN and DMN networks loaded almost exclusively on the Implicit and Explicit neural axes, respectively; Figure 4, left). However, as a noteworthy departure from this general observation, we found that only the frontoparietal control network exhibited substantial cross-loadings on the different neural axes. For example, Figure 5 shows the implicit/explicit loadings of individual regions within the Control subnetworks. Plotting the data in this way allows us to describe a region as being either *explicit-* or *implicit-aligned* if it has a greater loading on the Explicit or Implicit neural axis, respectively. Note that it is only the Control-A and Control-C networks that show meaningful alignment along this explicit-implicit dimension, with the posterior cingulate cortex (PCC) and precuneus regions in particular being strongly explicit-aligned.

**Figure 4:**
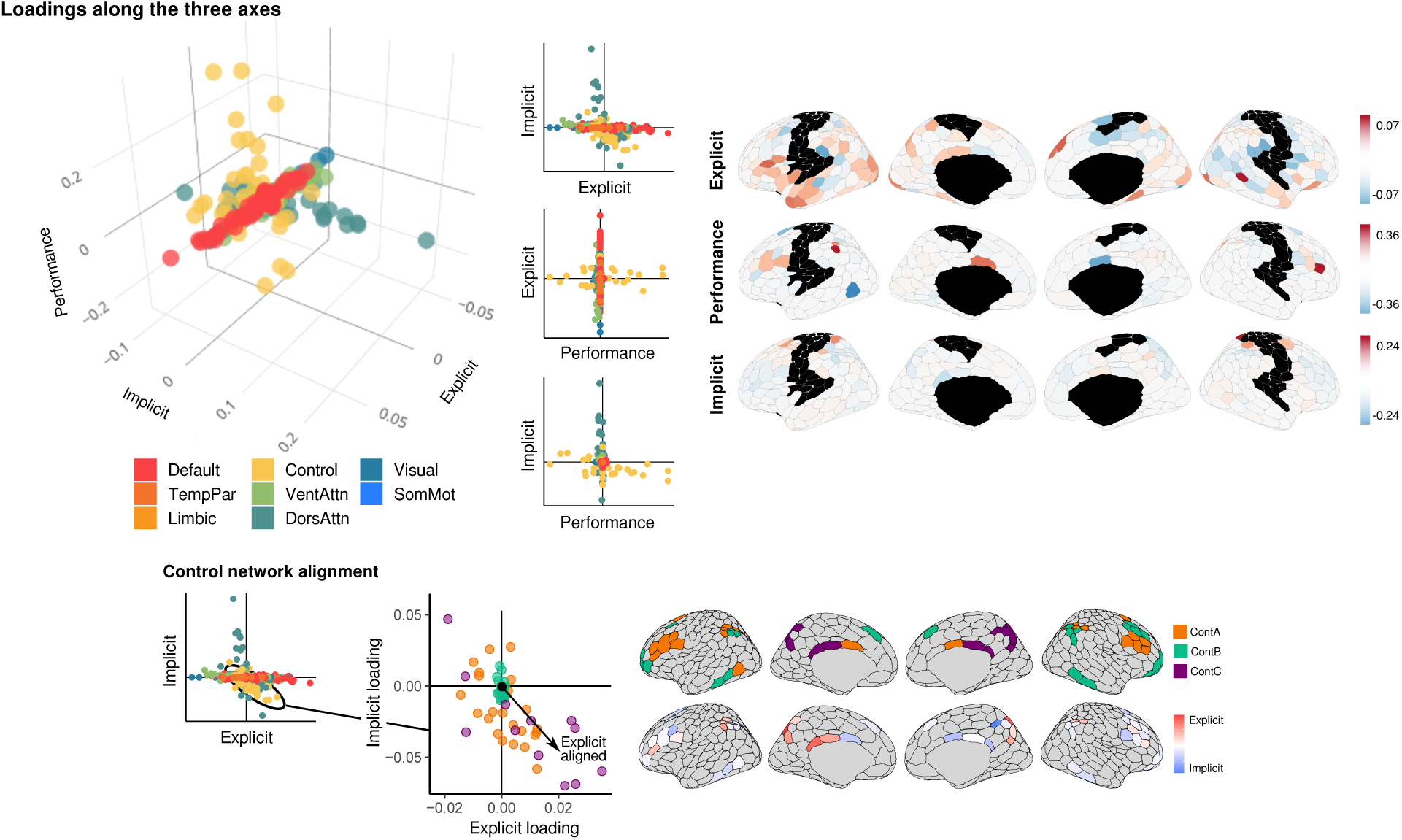
Alignment of individual cortical regions along the three separate neural axes – top. Left: Three-dimensional and two-dimensional projections show the loadings of each cortical parcel along each of the three neural axes (explicit, performance, and implicit). Right: Surface maps for each axis, with red/blue color scale denoting positive/negative coefficients. Note that regions in somatomotor cortex, with which all patterns of functional connectivity were assessed, are denoted in black. **Bottom)** Only regions of the control network showed meaningful cross-loadings between implicit and explicit axes. Of these subnetworks A and C were most variable, and the posterior cingulate and precuneus in particular were strongly associated with explicit learning.

**Figure 5:**
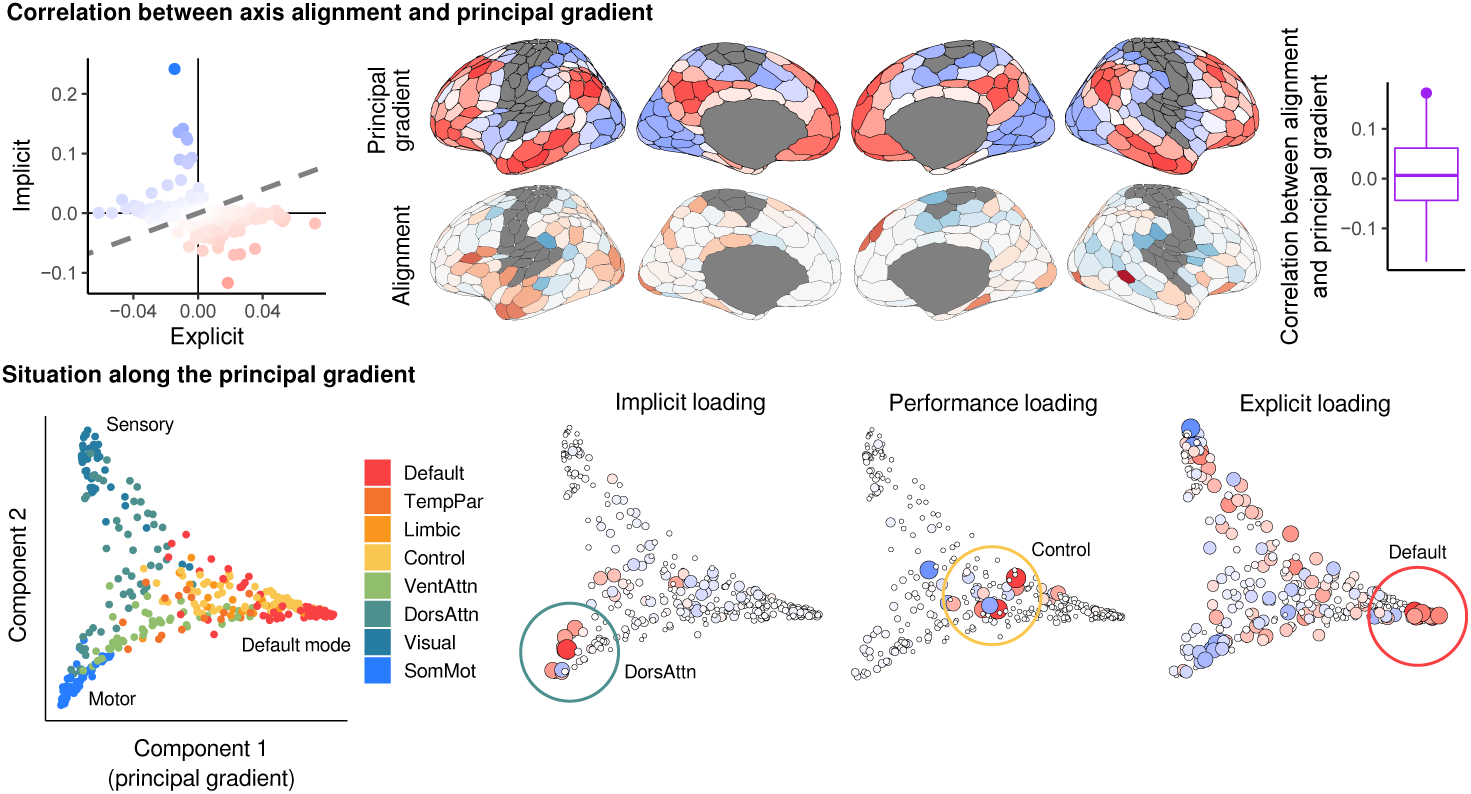
The learning-related neural axes correspond to the principal neural axis of brain organization – (Top) We characterized regions as being *explicit aligned* if they exhibited a higher loading on the explicit than on the implicit axis. We found that this *explicit alignment axis* was significantly correlated with the principal gradient [33]. Rightmost barplot indicates the null distribution of the correlation between the two brain maps as estimated by spin permutation testing. Note that whiskers denote the .025 and .975 quantiles of the distribution, while the point indicated the observed correlation. (**Bottom**) The three learning-related axes were compared with the principal cortical gradient characterized by [33] (see leftmost scatterplot). The implicit and explicit axes strongly feature regions at opposing ends of this principal gradient – closest to primary sensorimotor cortex (implicit), and closer to higher-order association cortex (explicit). Areas associated with the Performance neural axis tend to lie in between these opposing ends.

To more fully characterize the activity of each brain region in this way, we derived a single brain map – the explicit alignment axis – by computing the difference in rank for each ROI, such that positive values indicate relatively greater loading on the explicit, compared to implicit, neural axis, and vice versa for negative values (Figure 5; bottom). This single brain map captures the patterns of effects described above, with the greatest explicit alignment being observed in the DMN-B and Control-C subnetworks, and the least explicit (most implicit) alignment being observed in the superior parietal and premotor areas of the DAN-B subnetwork.

### 1.4 Learning-related neural axes correspond to features of functional cortical organization

The findings in our previous section suggest that explicit learning is supported by several subnetworks distributed across higher-order transmodal cortex whereas implicit learning is supported by more localized, sensorimotor areas within premotor and superior parietal cortex. To examine this idea more directly, we compared our explicit alignment neural axis to the well-known principal gradient of functional cortical organization that separates unimodal sensory and motor regions on one end, from higher-order transmodal regions belonging to the DMN and Control networks at the other end [33, see Figure 5]. When we tested for statistical significance (with Pearson’s r) using a spatial autocorrelation-preserving null model [“spin test”; 34], we found that our explicit alignment neural axis, derived from our motor learning study, was significantly positively correlated (p-spin < 0.05) with this principal functional cortical gradient, derived from spontaneous intrinsic activity. This suggests that the relative contribution of each brain region to explicit versus implicit learning processes roughly accords with its positioning along the unimodal-to-transmodal cortical hierarchy [33].

To more directly visualize this point, for each of our neural axes we have highlighted the loadings of each brain region along the principal axis of cortical functional organization (see Fig. 5, bottom). As can be seen in this figure, implicit and explicit aligned regions (denoted in red) tend to lie at opposite ends of the principal gradient, with implicit (resp. explicit) regions lying closer (resp. farther) to primary motor regions. We also find that areas with positive loadings in the Performance neural axis generally lie in between these implicit and explicit regions in the cortical hierarchy. Together, these results suggest that the whole-brain patterns of functional connectivity with motor cortex that we identify as being associated with different learning-related processes (i.e., implicit learning, task performance and explicit learning) can be partially explained as emergent features of a cortical processing hierarchy that extends from unimodal to transmodal cortex.

### 1.5 The explicit neural axis predicts rapid learning in a separate motor task

Implicit adaptation in the VMR task is thought to primarily reflect the updating – through sensory prediction errors – of an internal forward model for the sensory consequences of a motor command [5, 7]). In other motor learning tasks, where subjects are denied this sensory error feedback, this kind of updating is impossible, and so learning in such cases must rely on alternative neural systems [35, 36]. By contrast, explicit learning has been linked to various executive functions [37, 18, 38, 32], and the DMN – which features prominently in our Explicit neural axis – is suggested to specialize in cognitive computations that are independent of specific perceptual inputs [39]. We thus reasoned that the Explicit neural axis might generalize across sensorimotor learning tasks that provide different forms of sensory feedback, whereas the implicit neural axis would not.

To test this idea, in a different fMRI session we had our same participants perform a separate motor learning task in which they were required to alter their hand movements through purely reinforcement feedback. In this task, subjects used their right finger on the touchpad to trace – without visual feedback of their cursor – a rightward-curved path displayed on the screen (from a start position to target line). Following a baseline block (70 trials), in which subjects did not receive any feedback about their performance, they performed a learning block (200 trials). Here, they were instructed that they would now receive reward score feedback, between 0 and 100, based on how accurately they traced the visible path, and that they should attempt to maximize their score across trials. Critically -– and unbeknownst to the subjects — the reward score they received was actually based on how accurately they traced the (hidden) mirror-image path (reflected across the vertical axis; i.e., they would receive an imperfect score if they traced the visible path but would receive a 100 score if they perfectly traced the mirror image path; Figure 6, top left). Importantly, as the cursor was invisible, subjects could not use error-based learning mechanisms to improve their performance (as they could in the sensorimotor adaptation task). Rather, they could use only the scalar reward feedback, presented at the end of each trial, to refine their movements over time.

**Figure 6:**
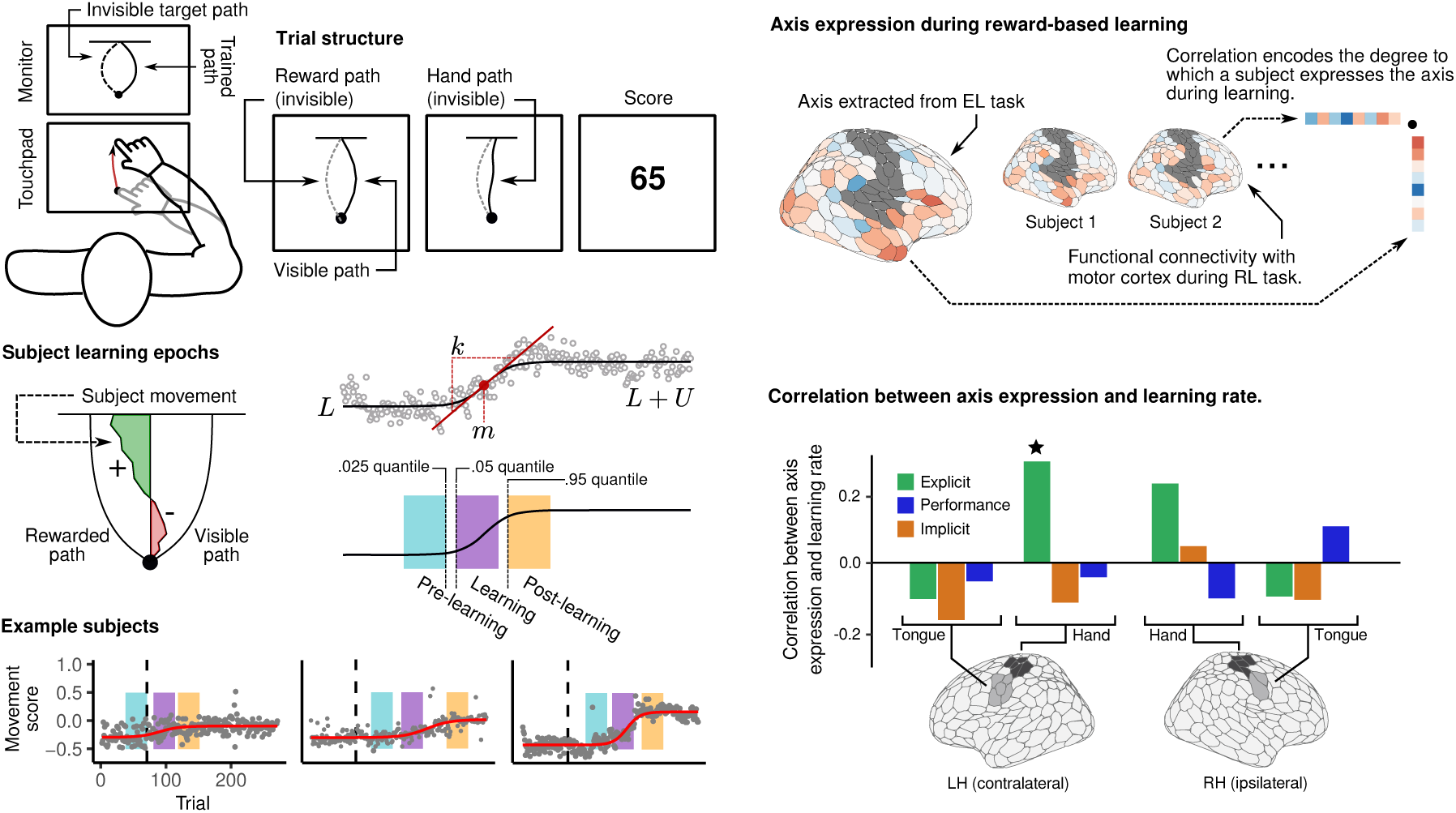
Subject performance and neural results for the separate reward-based motor learning task – (top left) Subjects learned to trace an invisible target trajectory with only score feedback. After learning to trace a rightward curved path (baseline trials), score feedback was introduced indicating (unbeknownst to the subject) their success in tracing the mirror image path (learning trials). **(Bottom left)** For each trial, we quantified the overall leftward or rightwardness of the response by taking the average x-position of the movement trajectory, with positive values indicating that the response was more towards the invisible target. We described this value as the *movement score*. Most subjects showed clearly defined periods of abrupt learning (which we termed “Aha! moments”), and we fit a sigmoid to each subjects’ movement scores in order to identify these periods. The .025, .05, and .95 quantiles of the resulting curve were used to define the end of the pre-learning, beginning of the learning, and beginning of the post-learning periods, respectively. **(Top right)** We examined whole-brain functional connectivity with motor cortex during the learning period, and asked whether learning-related increases in the expression of each neural axis was associated with faster learning. **(Bottom right)** Learning rate was significantly correlated with increases in the expression of the explicit – but not implicit or performance axes – during the abrupt learning period (see star). Notably, this effect was specific to patterns of functional connectivity with the contralateral hand area. When we re-derived the three axes using connectivity with either the ipsilateral hand area, or the contra- and ipsilateral tongue areas, this correlation with behaviour was abolished.

**Figure 7:**
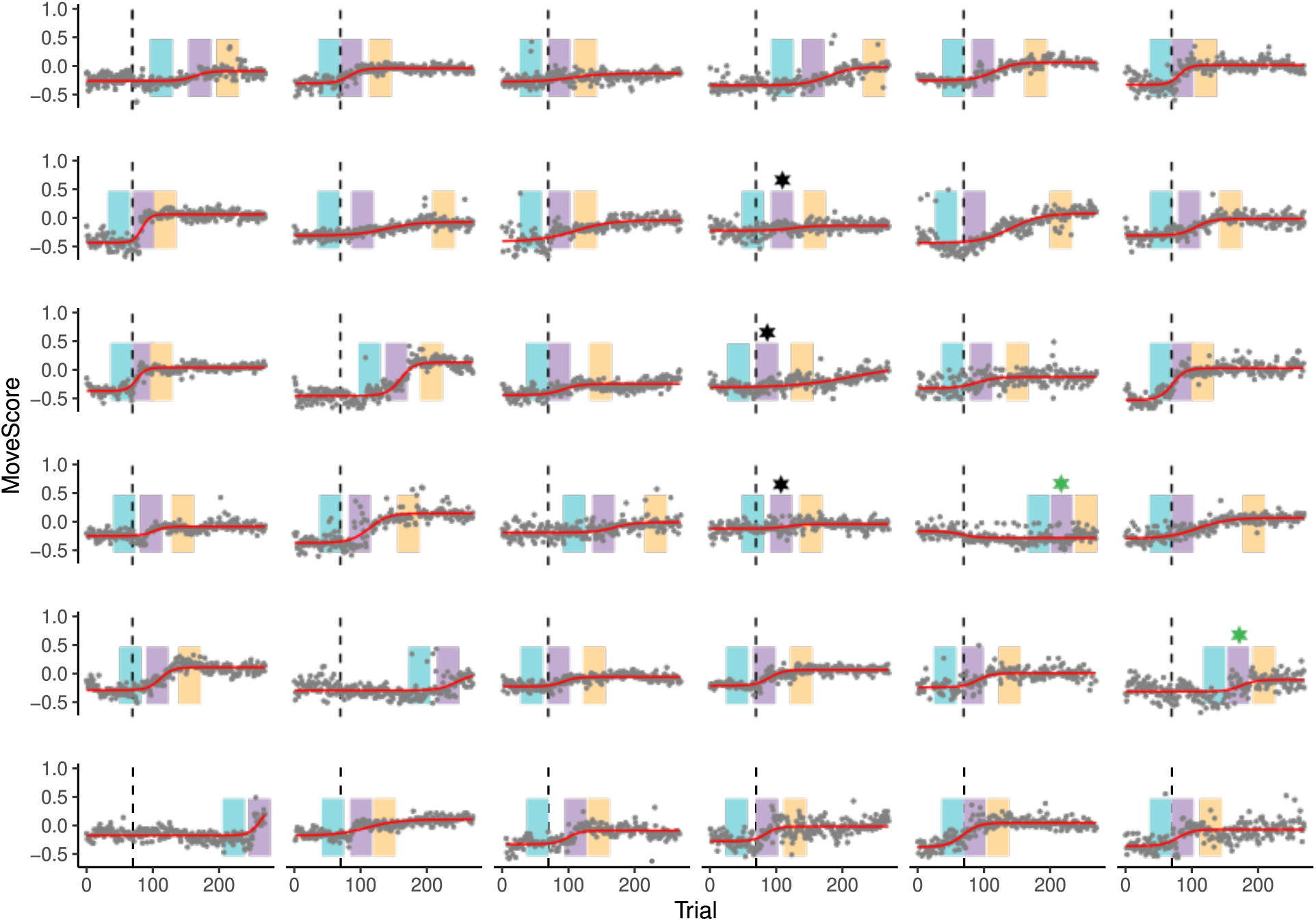
Reward-based learning task model fits for individual subjects –. Sigmoid fits (red lines) for individual subjects. Dots denote movement scores (mean horizontal cursor position) for individual trials, while dashed lines denote reward onset. Colored panels denote pre-learning, learning, and post-learning windows of 100 imaging volumes. Note that, in two subjects, learning periods occured so late that defining a post-learning period was impossible. Three subjects (black stars) exhibit performance patterns not well captured by a sigmoid function, and in particular generally exhibited flat or non-existent learning periods. For these subjects we nonetheless used the windows identified by the model fits, and simply recorded them as having very low learning rates. For two other subjects (green stars), we manually adjusted the windows in order to better capture the apparent learning onset.0

Learning under these types of conditions is known to be highly explicit [40]. Commensurate with this, we found that the majority of subjects displayed abrupt periods of rapid learning that tended to be book-ended by relatively flat performance (see example subjects in Figure 6). We termed these periods of abrupt learning as “Aha! moments”, as they clearly reflected participants’ sudden realization of the reward structure of the task (i.e., that it was not actually the visible path that was being rewarded). To identify these learning epoch, on each trial we computed a movement score quantifying the overall leftward (towards the invisible target) or rightwardness (towards the trained path) of the trajectory, and fit a sigmoid to each subjects’ performance. We defined pre-learning, learning, and post-learning periods based on the resulting fits (see Figure 6, left; see Methods and see supplement for fits to individual subjects). We then asked whether the expression of the Explicit neural axis during the learning period was related to subjects’ learning rates. Specifically, for each subject, we computed the spatial correlation between each of our learning-related neural axes extracted from the VMR task (i.e., the Implicit, Performance, and Explicit axes) with the observed patterns of functional connectivity during each of the periods (pre-learning, learning, and post-learning) associated with the reward-based learning task. Note that these spatial correlations provide us with an index of the degree to which each neural axis from the sensorimotor adaptation task is expressed during the different periods of reward-based learning, and we thus call these correlation the *explicit* (resp. performance, implicit) axis scores. Next, we examined whether these axis scores were correlated with subjects’ learning rates.

The results of this analysis are shown in Figure 6 (bottom right). Consistent with our predictions, we observed a significant positive correlation between the expression of the Explicit neural axis (derived from the VMR task) during the learning period of the reward-based task, with subjects’ learning rates. That is, the more similar the patterns of functional connectivity during the rapid learning period of the reward-task was to the patterns of functional connectivity associated with the Explicit neural axis, the more rapidly subjects learned the hidden path (*r* = .34*, t*_34_ = 2.04*, p* = .049).

Importantly, as a control analysis, we observed no such relationship for either of the Implicit (*r* = *−.*12*, t*_34_ = *−.*71*, p* = .48) or Performance (*r* = *−.*05*, t*_34_ = *−.*29*, p* = .77) neural axes (Figure 6, bottom right). Note that the correlation of the explicit axis with performance was also abolished when we examined the post-learning period of RL task performance (*r* = *−*0.06, *t*_34_ = *−*0.34, *p* = 0.74).

To further examine the specificity and functional relevance of these effects, we re-derived each of our neural axes during the VMR task (Implicit, Performance, and Explicit) using whole-brain functional connectivity with the *ipsilateral* hand area of the motor cortex, as well as the primary motor tongue area in each hemisphere (Figure 6, bottom right). Critically, in none of these cases did we observe a significant correlation between the expression of these newly derived Explicit neural axes and subjects’ learning rates. Note that, although the correlation between Explicit axis expression and learning rate was not significant in the right hand area, neither was the difference in correlation between the two hemispheres significant.

Taken together, these findings indicate that it is the expression of the Explicit neural axis, and not the Implicit or Performance neural axes, that predicts abrupt learning in a separate motor task that does not afford learning via sensory prediction errors. Moreover, our analyses demonstrate the selectivity of this effect; we find that these results are exclusive to (1) neural activity during the abrupt learning phase (and not pre- and post-learning phases) and, (2) connectivity with the *contralateral* hand area of motor cortex, indicating that these patterns of connectivity are neuroanatomically and behaviourally relevant.

## 2 Discussion

Adaptive motor behaviour is achieved through a combination of distinct learning processes, yet the specific functional interactions between brain regions that support these processes are poorly understood. Here we examined learning-related changes in human cortical, striatal and cerebellar functional connectivity with the primary motor cortex while individuals performed a classic sensorimotor adaptation task. Using subjects’ explicit reports, as well as their visuomotor errors experienced during adaptation, we isolated three distinct axes of connectivity during early learning that corresponded to subjects’ use of implicit and explicit learning processes, as well as their learning performance. Whereas we found that implicit learning correlated with connectivity changes between motor cortex and superior parietal and premotor regions (the Implicit neural axis), we found that explicit learning correlated with connectivity changes between motor cortex and higher-order transmodal cortex, in particular the DMN (the Explicit neural axis). Notably, we distinguished both of these connectivity changes from activity patterns related subjects’ performance errors during learning, which we found was correlated with the activity of the frontoparietal control network (the Performance neural axis). Further analyses showed that these three distinct neural axes could be mapped onto different levels of the cortical hierarchy, which extends from primary sensory and motor systems to higher-order transmodal cortex. Finally, we examined the extent to which these neural axes generalized to a separate motor task known to involve explicit learning processes, and found that it was the expression of the Explicit neural axis–and not the Implicit and Performance axes–that predicted subject learning under these circumstances.

The main contribution of our study, therefore, was in establishing a tripartite distinction in terms of how connectivity between motor cortex and other regions of the brain relate to implicit and explicit learning, as well as processes more directly linked to performance. Whereas in behavioural studies of motor learning these different factors can be readily differentiated and measured [13, 7, 22], to date it has been difficult to establish the neural bases of these factors. For instance, while activation changes observed during early learning may reflect subjects’ use of cognitive strategies, as is often interpreted [27], these changes may also reflect modulations in sensory feedback associated with the use of such strategies (e.g., decreased visual errors), or even changes related to trial-by-trial recalibration of the internal model (which occur automatically in parallel with the use of strategies)[41, 13, 42, 7]. Our study overcame several of these limitations by obtaining a separate measure of subjects’ actual re-aiming strategy, and by controlling for performance-related effects on our measures subjects’ explicit and implicit learning. This allowed us to identify unique neural signatures of these distinct processes in patterns of functional connectivity between motor cortex and the rest of the brain.

One important feature of our results is our corroboration of prior suggestions that superior parietal and premotor areas of the DAN are important for implicit processes during motor learning [14, 15, 16]. Contemporary theories of sensorimotor adaptation, based on optimal feedback control, highlight these regions along with the cerebellum in the maintenance and updating of an internal forward model in response to sensory feedback [43, 44]. Although we found that the cerebellum loaded relatively weakly onto our Implicit neural axis, we note that its anterior motor regions, particularly in the right (ipsilateral) cerebellum, were aligned to this axis. This is consistent with the neuroanatomy and known role of this brain structure in computing the sensory prediction errors that drive implicit motor adaptation [45, 46, 36, 47, 48].

At the same time, our study shows for the first time that explicit processes during learning are associated with functional connectivity between motor cortex and regions of the DMN, as well as Control network areas in the PCC and precuneus. Prior work has often presumed that explicit learning is mainly associated with connectivity changes in “task-positive” frontoparietal brain areas, such as the DLPFC [10, 12, 27, 31]. Indeed, these areas tend to activate across a wide variety of task conditions requiring executive processes, such as spatial working memory, which is thought to support explicit re-aiming strategies during VMR learning [49, 37, 38, 50, 51]. Consequently, increases in DLPFC activity and connectivity during early learning, and its relationship to subject performance [37, 27], has often been interpreted to reflect working memory processes related to the implementation of an explicit re-aiming strategy. However, in the current study we demonstrate that functional connectivity with the DLPFC (Control-A subnetwork) is linked not directly with explicit learning itself, but rather with improvements in subject visuomotor performance (error reduction, see also [52]). This suggests that the DLPFC’s role during motor learning aligns more closely with its established functions in target selection and performance monitoring [53, 54, 55]. In this regard, it is noteworthy that the DLPFC and cingulate cortex – also grouped under the same Control A subnetwork – has more generally been proposed to support the task-dependent updating of behaviour in response to errors [55].

As for the DMN and PCC/precuneus cortical regions which featured prominently in the Explicit neural axis, it is important to note that prior sensorimotor adaptation studies have not commonly focused on the activity in these higher-order brain areas. This is perhaps not surprising, given the assumption that these regions are primarily “task-negative” (i.e., tend to be suppressed during tasks; [39]). However, outside the domain of motor control, the role of the DMN in explicit processes has been well-documented. The DMN has been implicated in multiple states such as self-referential processing, theory of mind, and spontaneous states such as mind wandering – all of which depend on explicit conscious attention [17, 56, 57, 58, 59, 39]. Moreover, there is also emerging neurophysiological evidence that DMN areas are uniquely engaged in conditions when performance is initially poor and when executive control is advantageous [60, 61], such as during early learning. For instance, work in nonhuman primates has shown that activity in DMN areas is linked to action exploration and the use of strategies during task performance [62]. This suggests a key role for DMN areas in the organization of cognitive resources when performance errors are initially high [61, 63, 62].

In this way, our study suggests that, like in multiple other domains, the DMN may play a role in motor control that is linked to the capacity of humans to explicitly attend to their own thoughts and actions – a skill that is particularly important when behaviour must be guided by by internally generated strategies and rule-sets, as well as information accrued over longer time horizons than by in-the-moment sensory inputs [39]. This contribution of the DMN to behavior is hypothesized to be enabled through it’s unique topographic positioning on cortex [33], with each core node of the DMN being located maximally distant from sensory and motor systems. This positioning is thought to free DMN areas from the constraints of extrinsically driven neural activity, allowing these regions to play a role in the orchestration of information processing across distributed functional systems [64, 39]. Consistent with these properties, we showed that our explicit alignment axis was significantly positively correlated with the principal cortical axis identified by [33], spanning from unimodal to transmodal cortex. As noted by others [65], this cortical architecture closely resembles functional differences in the time horizons over which sensorimotor and transmodal areas are known to process information. For instance, previous research has demonstrated that the temporal windows of information processing change in a hierarchical fashion across cortex [66, 67, 68], with sensorimotor areas integrating information over the order of milliseconds to seconds versus transmodal areas that integrate information over the order of seconds to minutes and longer. These longer “temporal receptive windows” in transmodal cortex [66, 67] are thought to enable these areas to encode more slowly changing states of the world and incorporate acquired knowledge into more complex scenarios -– attributes that are both crucial when searching for and implementing cognitive strategies during learning.

A final unique feature of the current study was in exploring the extent to which the neural axes identified during sensorimotor adaptation generalized across other forms of motor learning. Although few neural studies have examined the use of reinforcement mechanisms during motor learning, there is considerable behavioural evidence indicating that explicit processes are particularly crucial in tasks that deny the kinds of sensory feedback that drive error-based adaptation, such as in our reward-based motor task, or the tasks of [19, 69, 70, 71]. In these types of reward-based motor learning tasks, the agent is provided no feedback indicating precisely how, or to what extent, the executed versus optimal movements differ, and so they must perform some kind of credit assignment in order to decide which feature of the movement to adjust on subsequent trials [72]. In these contexts, subjects will often develop an explicit understanding of the task structure, with their motor responses demonstrating a clear prioritization of the features relevant to performance [70, 69]. Consistent with this, the majority of subjects in our reward-based motor task exhibited periods of abrupt learning (which we called ’Aha! moments’), suggesting a sudden realization of the correct movement trajectory. These abrupt shifts in behaviour have been studied in the context of one-shot learning, where research has implicated regions of the DMN [73] in facilitating rapid learning through an explicit record of past action-outcome associations [74]. As noted above, these regions featured prominently in our Explicit neural axis, the expression of which predicted subject performance in the reward-based motor task.

Although our study identifies distinct patterns of connectivity with motor cortex during sensorimotor adaptation and the component processes that these patterns support, it also leaves several important questions for future research. First, because we restricted our analysis to bipartite functional interactions between the motor cortex and the rest of the brain, we cannot speak to connectivity changes occurring *outside* of the motor cortex and how they may relate to these different learning processes. For instance, our analysis does not preclude communication between regions of visual cortex and frontoparietal regions in the monitoring of visuomotor performance. Nevertheless, we believe our study provides a critical first step in understanding how interactions with the motor system drive sensorimotor adaptation. Second, although we included regions of both the striatum and cerebellum in our analyses, we found that they tended to load relatively weakly on the neural axes as compared to cortical regions. This is likely to reflect relatively stronger cortico-cortical functional connectivity (compared to cortico-extracortical connectivity), rather than the non-involvement of these regions in learning. Indeed, as cortico-cerebellar/striatal functional connectivity is relatively weaker, it may simply be less predictively useful in our regression analyses than using cortico-cortical connectivity. Future work could construct models based solely on cortico-extracortical connectivity to more precisely identify the role of extracortical regions in different learning processes. It may also be important to examine the role of these regions using high-field strength fMRI, as it may have the spatial fidelity necessary to unveil the complex role that these regions play in coordinating cortex-wide interactions.

## 3 Methods

For detailed behavioural and neuroimaging methods, see Supplemental Methods.

## Acknowledegments

C.N.A. was supported by a Natural Sciences and Engineering Research Council (NSERC) graduate award. J.P.G. was also supported by a NSERC Discovery Grant (RGPIN-2017-04684), Canadian Institutes of Health Research Grant (MOP126158), and Botterell Foundation Award, as well as funding from the Canadian Foundation for Innovation (35559). The authors would like to thank Martin York and Sean Hickman for technical assistance, and thank Don O’Brien for assistance with data collection.

## Data availability

Behavioral and imaging data are archived at Queens University and are available from the authors upon reasonable request.

## Code availability

Imaging data were preprocessed using fmriPrep 1.4.0, which is open source and freely available. Operations on covariance matrices, including estimation and centering, were performed using the R package spdm, which is freely available in a repository at https://github.com/areshenk-rpackages/spdm. Tutorial code and data for implementing the centering procedure and gradient analyses are hosted in a GitHub repository at https://github.com/areshenk-opendata/2023-implicitexplicit.

### 3.1 Participants

Forty-six right-handed subjects (27 females, aged 18-28 years, mean age: 20.3 years) participated in three separate testing sessions, each spaced approximately 1 week apart: the first, an MRI-training session, was followed by two subsequent MRI experimental sessions. Of these 46 subjects, 10 participants were excluded from the final analysis [1 subject was excluded due of excessive head motion in the MRI scanner (motion greater than 2 mm or 2° rotation within a single scan); 1 subject was excluded due interruption of the scan during the learning phase of the reward-based motor task; 5 subjects were excluded due to poor behavioural performance in one or both tasks (4 of these participants were excluded because >25% of trials were not completed within the maximum trial duration; and one because >20% of trials had missing data due to insufficient pressure of the fingertip on the MRI-compatible tablet); 3 subjects were excluded due to a failure to properly perform the reward-based task (these subjects did not trace the visible rightward path during the baseline phase, but rather continued to trace a straight line for the entire duration of the task, suggesting that they did not attend to the task)].

Right-handedness was assessed with the Edinburgh handedness questionnaire [75]. Participants’ informed consent was obtained before beginning the experimental protocol. The Queen’s University Research Ethics Board approved the study and it was conducted in coherence to the principles outlines in the Canadian Tri-Council Policy Statement on Ethical Conduct for Research Involving Humans and the principles of the Deceleration of Helsinki (1964).

### 3.2 Procedure

In the first session, participants took part in an MRI-training session inside a mock (0 T) scanner, that was made to look and sound like a real MRI scanner. We undertook this training session to (1) introduce participants to features of the VMR and reward-based motor tasks that would be subsequently performed in the MRI scanner, (2) ensure that subjects obtained baseline performance levels on those two tasks, and (3) ensure that participants could remain still for 1.5 hr. To equate baseline performance levels across participants, subjects performed 80 trials per task. To train participants to remain still in the scanner, we monitored subjects’ head movement via a Polhemus sensor attached to each subject’s forehead (Polhemus, Colchester, Vermont). This allowed a real-time read-out of subject head displacement in the three axes of translation and rotation (6 dimensions total). Whenever subjects’ head translation and/or rotation reached 0.5 mm or 0.5° rotation, subjects received an unpleasant auditory tone, delivered through a speaker system located near the head. All of the subjects used in the study learned to constrain their head movement through this training regimen. Following this first session, subjects then subsequently participated in reward-based VMR motor tasks, respectively (see below for details).

### 3.3 Apparatus

During testing in the mock (0 T) scanner, subjects performed hand movements that were directed towards a target by applying finger tip pressure on a digitizing touchscreen tablet (Wacom Intuos Pro M tablet). During the actual MRI testing sessions, subjects used an MRI-compatible digitizing tablet (Hybridmojo LLC, CA, USA). In both the mock and real MRI scanner, the target and cursor stimuli were rear-projected with an LCD projector (NEC LT265 DLP projector, 1024 x 768 resolution, 60 Hz refresh rate) onto a screen mounted behind the participant. The stimuli on the screen were viewed through a mirror fixated on the MRI coil directly above the participants’ eyes, thus preventing the participant from being able to see their hand.

### 3.4 Sensorimotor adaptation task

To study sensorimotor adaptation, we used the well-characterized visuomotor rotation (VMR) paradigm [21, ; Figure **??**, top left]. During the VMR task, participants performed baseline trials in which they used their right index finger to perform center-out target-directed movements. After these baseline trials, we applied a 45*^◦^* clockwise (CW) rotation to the viewed cursor, allowing investigation of VMR learning. Following this, we assessed participants’ re-aiming strategy associated with their learning.

Each trial started with the participant moving the cursor (3 mm radius cyan circle) into the start position (4 mm radius white circle) in the centre of the screen by sliding the index finger on the tablet. To guide the cursor to the start position, a ring centred around the start position indicated the distance between the cursor and the start position. The cursor became visible when its centre was within 8 mm of the centre of the start position. After the cursor was held within the start position for 0.5 s, a target (5 mm radius red circle) was shown on top of a grey ring with a radius of 60 mm (i.e., the target distance) centred around the start position. The target was presented at one of eight locations, separated by 45*^◦^* (0, 45, 90, 135, 180, 225, 270 and 315*^◦^*), in pseudorandomized bins of eight trials. Participants were instructed to hit the target with the cursor by making a fast finger movement on the tablet. They were instructed to ‘slice’ the cursor through the target to minimize online corrections during the reach. If the movement was initiated (i.e., the cursor had moved fully out of the start circle) before the target appeared, the trial was aborted and a feedback text message “Too early” appeared centrally on the screen. In trials with correct timing, the cursor was visible during the movement to the ring and then became stationary for one second when it reached the ring, providing the participant with visual feedback of their endpoint reach error. If any part of the stationary cursor overlapped with any part of the target, the target coloured green to indicate a hit. Each trial was terminated after 4.5 s, independent of whether the cursor had reached the target. After a delay of 1.5 s, allowing time to save the data, the next trial started with the presentation of the start position.

During the participant training session in the mock MRI scanner (i.e., 2-weeks prior to the VMR MRI testing session), participants performed a practice block of 40 trials with veridical feedback (i.e., no rotation was applied to the cursor). This training session exposed participants to several key features of the task (e.g., use of the touchscreen tablet, trial timing, presence and removal of cursor feedback), and allowed us to establish adequate performance levels.

At the beginning of the MRI testing session, but prior to the first scan being collected, participants re-acquainted themselves with the VMR task by performing 80 practice trials with veridical cursor feedback. Next, we collected an anatomical scan, followed by the baseline fMRI experimental run, wherein subjects performed 64 trials of the VMR task with veridical cursor feedback using their right hand. Next, they performed the learning scan, wherein subjects performed 160 trials in which feedback of the cursor during the reach was rotated clockwise by 45*^◦^*. Following this fMRI experimental run, participants were asked to report their strategic aiming direction (over 16 trials). In these trials, a line between the start and target position appeared on the screen at the start of each trial. Participants were asked to use a separate MRI joystick (Current Designs, Inc.) positioned at their left hip to rotate the line to the direction that they would aim their finger movement in for the cursor to hit the target, and click the button on the joystick box when satisfied. Following the button click, the trial proceeded as a normal reach trial. These 16 “report” trials were followed by 16 normal rotation trials. As several subjects expressed confusion about the nature of the report trials, often failing to adjust the line at all during the first few trials, we discarded the first 8 report trials for all subjects, and calculated the mean aim direction using the final 8 trials. Note that we had subjects perform these report trials following learning [as in 69] given our prior behavioural work showing that the inter-mixing of report trials during learning can lead to more participants adopting an explicit, re-aiming strategy [22, 69], thereby distorting participants’ learning curves. Participants were not informed about nature or presence of the visuomotor rotation before or during the experiment.

Note that all subjects performed the VMR task after having performed the reward-based motor task (i.e., testing order was not counterbalanced across the RL and EL tasks). We made this decision in light of our knowledge that subjects may alter their behavior after performing explicit report trials in the VMR task [22] (i.e., often developing explicit awareness of the perturbation and using that to guide their performance). We were concerned that, had subjects performed the VMR task first, they may have anticipated some experimental manipulation (or deception), and thus would not have approached the reward-based motor task in a naive fashion. From this perspective, our approach of having defined the explicit and implicit neural axes on the (second) VMR task and then using those neural axes to predict performance on the (first) reward-based motor task, strengthens our interpretations of the effects.

### 3.5 Reward-based Learning task

In the reward-based motor task (Figure 6, left), participants learned to produce, through reward-based feedback, finger movement trajectories with a specific (unseen) shape. Specifically, subjects were instructed to repeatedly trace, without visual cursor feedback of their actual finger paths, a subtly curved path displayed on the screen (the visible path). During learning trials, participants were told that, following each trial, they would receive a score based on how accurately they traced the visible path (and were instructed to maximize points across trials). However, unbeknownst to them, they actually received points based on how well they traced the mirror-image path (the target path). Critically, because participants received no visual feedback about their actual finger trajectories or the ‘rewarded’ shape, they could not use error-based information to guide learning. Our task was inspired by the motor learning tasks developed by [71, 70].

Each trial started with the participant moving the cursor (3 mm radius cyan circle) into the start position (4 mm radius white circle) at the bottom of the screen by sliding their index finger on the tablet. The cursor was only visible when it was within 30 mm of the start position. After the cursor was held within the start position for 0.5 s, the cursor disappeared and a curved path and a target distance marker appeared on the screen. The target distance marker was a horizontal red line (30 x 1 mm) that appeared 60 mm above the start position. The visible path connected the start position and target distance marker and had the shape of a half sine wave with an amplitude of 0.15 times the target distance. Participants were instructed to trace the curved visible path. When the cursor reached the target marker distance, the marker changed colour from red to green to indicate that the trial was completed. Importantly, participants did not receive feedback about the position of their cursor during the trial.

In the baseline block, participants did not receive feedback about their performance. In the learning block, participants were rewarded 0 to 100 points after reaching the target distance marker, and participants were instructed to do their best to maximize this score across trials (the points were displayed as text centrally on the screen). They were told that to increase the score, they had to “trace the line more accurately”. Each trial was terminated after 4.5 s, independent of whether the cursor had reached the target. After a delay of 1.5 s, allowing time to save the data, the next trial started with the presentation of the start position.

To calculate the score on each trial in the learning block, the x position of the cursor was interpolated at each cm displacement from the start position in the y direction (i.e., at exactly 10, 20, 30, 40, 50 and 60 mm). For each of the six y positions, the absolute distance between the interpolated x position of the cursor and the x position of the rewarded path (mirror image of visible path) was calculated. The sum of these errors was scaled by dividing it by the sum of errors obtained for a half cycle sine-shaped path with an amplitude of 0.5 times the target distance, and then multiplied by 100 to obtain a score ranging between 0 and 100. The scaling worked out so that a perfectly traced visible path would result in an imperfect score of 40 points. This scaling was chosen on the basis of extensive pilot testing in order to achieve variation in subject performance, and to ensure that subjects still received informative score feedback when tracing in the vicinity of the visible trajectory.

During the participant training session in the mock MRI scanner (i.e., 1-week prior to the MRI testing session), participants only performed a practice block in which they traced a straight line with (40 trials) and then without (40 trials) visual feedback of the position of the cursor during the reach. As with the VMR task, this training session exposed participants to several key features of the task (e.g., use of the touchscreen tablet, trial timing, used of cursor feedback to correct for errors) and allowed us to establish adequate baseline performance levels. Importantly, however, this training session did not allow for any reward-based learning to take place.

At the beginning of the MRI testing session, but prior to the first scan being collected, participants re-acquainted themselves with the reward-based motor task by first performing a practice block in which they traced a straight line with (40 trials) and then without (40 trials) visual feedback of the position of the cursor during the reach. Next, we collected an anatomical scan, following by a DTI scan, followed by a resting-state fMRI scan. During the latter resting-state scan, participants were instructed to rest with their eyes open while fixating on a central cross location presented on the screen. Following this, participants performed the reward-based motor task, which consisted of two separate experimental runs without visual feedback of the cursor: (1) a baseline block of 70 trials in which they attempted to trace the curved visible path and no score was provided, and (2) a separate learning block of 200 trials in which participants were instructed to maximize their score shown at the end of each trial.

### 3.6 Behavioral data analysis

#### 3.6.1 VMR task analysis

Trials in which the reach was initiated before the target appeared ( 4% of trials) or in which the cursor did not reach the target within the time limit ( 5% of trials) were excluded from the offline analysis of hand movements. As insufficient pressure on the touchpad resulted in a default state in which the cursor was reported as lying in the top left corner of the screen, we also excluded trials in which the cursor jumped to this position before reaching the target region ( 2% of trials). We then applied a conservative threshold on the movement and reaction times, removing the top .05% of trials across all subjects. As the VMR task required the subject to determine the target location prior to responding, we also set a lower threshold of 100 ms on the reaction time.

We then calculated the angle difference between the target position and the last sample before the cursor exceeded the target circle. This was then converted to an adaptation score by subtracting the angle of the rotation on each trial (0*^◦^* during baseline, and 45*^◦^* during learning) in order to derive the actual aim direction on each trial.

#### 3.6.2 Reward-based motor task analysis

Trials in which the cursor did not reach the target distance marker within the time limit were excluded from the offline analysis of hand movements ( 1% of trials). As in the VMR task, we excluded trials in which insufficient pressure was applied ( 2% of trials), and also separately applied a conservative threshold on the movement and reaction times, removing the top .05% of trials across all subjects. As the VMR task did not involve response discrimination, we did not set a lower threshold on these variables.

For each trial, we computed a *movement score* by integrating the horizontal position of the trajectory, so as to derive a scalar measure of the overall leftward or rightwardness of the movement. Note that we flipped the sign of this measure so that positive (resp. negative) values denote trajectories closer to the target (resp. trained) trajectory. To each subjects’ movement data, we fit a generalized sigmoid function in which the predicted movement score on each trial *x* was given by

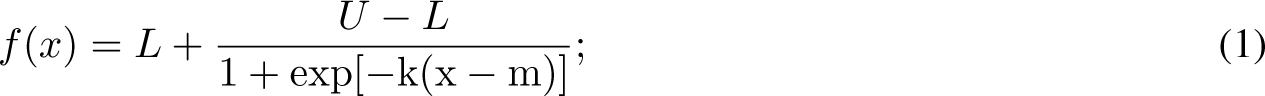

where *L* and *U* denote the lower and upper asymptotes, respectively; *m* denotes the midpoint of the learning period; and *k* denotes the learning rate (but see our comments below). Because the fits were unstable for several subjects with very slow learning rates, we regularized the fits with the following Bayesian model:

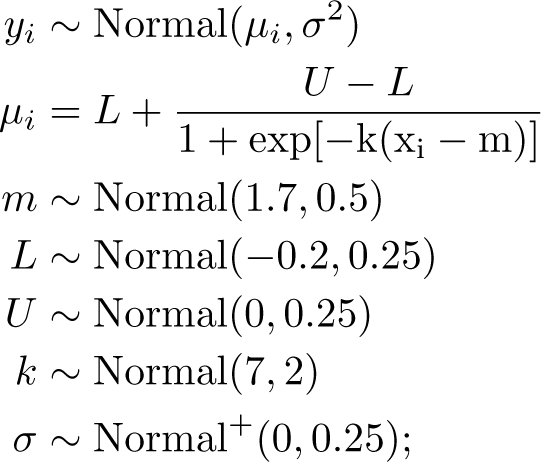

Note that trial numbers were scaled by 100 to keep the units comparable, so that the prior mean of 1.7 on the midpoint *m* was centered at trial 170 (i.e. the center of the learning block). The prior mean of *−*0.25 on *L* was centered about the movement score corresponding to tracing the initial learned trajectory, while the prior mean on *U* corresponds to the midpoint between perfect learning and an absence of learning. The prior scale of 0.25 on the error standard deviation was chosen empirically to place high probability mass over the standard deviations observed in subjects’ baseline blocks.

Models were fit using the Hamiltonian Monte-Carlo algorithm implemented by the Stan programming language [76]. For each subject, we fit the model using three chains of 5000 samples, of which the first 1500 were discarded as burn-in. In each case, visual inspection of the chains suggested adequate convergence, with potential scale reduction factor *R*^^^ less than 1.01 for all parameters and all subjects.

For each subject, pre-learning and learning epochs were defined using the posterior mean parameter estimates. Specifically, we defined the end of the pre-learning epoch to be the point at which the posterior mean curve exceeded 2.5% of its total range *U − L*. The beginning of the learning epoch was then defined to be the point at which the curve exceeded 5% of its total range. Using these boundaries, we then defined pre-learning and learning periods of 100 imaging volumes. Several example subjects are shown in Figure 6 (bottom left), and we have included illustrations of the model fits for every subject as supplemental material.

Visual inspection of the model fits indicated that the sigmoid generally accurately captured the overall pattern of subjects’ learning, with a few exceptions. In the supplement, we note three subjects who did not display well defined learning periods, and for whom the fitted sigmoid was nearly flat. In these cases, we retained the identified pre-learning and learning epochs, and these subjects were simply recorded as having extremely low learning rates. Two subjects were poorly characterized by the sigmoid fit, and so we manually identified pre-learning and learning epochs in these cases. Each of these subjects is indicated in the supplemental figure. In these latter two subjects, the parameter *k* did not encode the learning rate within the learning epoch itself, and so for each subject we defined their learning rate (for the analysis reported in Figure 6) to be the slope of the line of best fit to the scores in their learning epoch.

### 3.7 MRI Acquisition

Participants were scanned using a 3-Tesla Siemens TIM MAGNETOM Trio MRI scanner located at the Centre for Neuroscience Studies, Queen’s University (Kingston, Ontario, Canada). Functional MRI volumes were acquired using a 32-channel head coil and a T2*-weighted single-shot gradient-echo echo-planar imaging (EPI) acquisition sequence (time to repetition (TR) = 2000 ms, slice thickness = 4 mm, in-plane resolution = 3 mm x 3 mm, time to echo (TE) = 30 ms, field of view = 240 mm x 240 mm, matrix size = 80 x 80, flip angle = 90°, and acceleration factor (integrated parallel acquisition technologies, iPAT) = 2 with generalized auto-calibrating partially parallel acquisitions (GRAPPA) reconstruction. Each volume comprised 34 contiguous (no gap) oblique slices acquired at a 30° caudal tilt with respect to the plane of the anterior and posterior commissure (AC-PC), providing whole-brain coverage of the cerebrum and cerebellum. Each of the task-related scans included an additional 8 imaging volumes at both the beginning and end of the scan. On average, each of the MRI testing sessions lasted 2 hrs.

At the beginning of the reward-based motor task MRI testing session, a T1-weighted ADNI MPRAGE anatomical was also collected (TR = 1760 ms, TE = 2.98 ms, field of view = 192 mm x 240 mm x 256 mm, matrix size = 192 x 240 x 256, flip angle = 9°, 1 mm isotropic voxels). This was followed by a series of Diffusion-Weighted scans, wherein we acquired two sets of whole-brain diffusion-weighted volumes (30 directions, b = 1000 s mm-2, 65 slices, voxel size = 2 x 2 x 2 mm3, TR = 9.3 s, TE = 94 ms) plus 2 volumes without diffusion-weighting (b = 0 s mm-2). Next, we collected a resting-state scan, wherein 300 imaging volumes were acquired. For the baseline and learning scans during the motor task, 222 and 612 imaging volumes were acquired, respectively.

At the beginning of the VMR MRI testing session, we gathered high-resolution whole-brain T1-weighted (T1w) and T2-weighted (T2w) anatomical images (in-plane resolution 0.7 x 0.7 mm2; 320 x 320 matrix; slice thickness: 0.7 mm; 256 AC-PC transverse slices; anterior-to-posterior encoding; 2 x acceleration factor; T1w TR 2400 ms; TE 2.13 ms; flip angle 8°; echo spacing 6.5 ms; T2w TR 3200 ms; TE 567 ms; variable flip angle; echo spacing 3.74 ms). These protocols were selected on the basis of protocol optimizations designed by [77]. Following this, for the baseline and learning functional scans, 204 and 492 imaging volumes were acquired, respectively.

### 3.8 fMRI Preprocessing

Results included in this manuscript come from preprocessing performed using *fMRIPrep* 1.4.1 ([78]; [79]; RRID:SCR_016216), which is based on *Nipype* 1.2.0 ([80]; [81]; RRID:SCR_002502).

#### Anatomical data preprocessing

A total of 2 T1-weighted (T1w) images were found within the input BIDS dataset. All of them were corrected for intensity non-uniformity (INU) with N4BiasFieldCorrection [82], distributed with ANTs 2.2.0 [83, RRID:SCR_004757]. The T1w-reference was then skull-stripped with a *Nipype* implementation of the antsBrainExtraction.sh workflow (from ANTs), using OASIS30ANTs as target template. Brain tissue segmentation of cerebrospinal fluid (CSF), white-matter (WM) and gray-matter (GM) was performed on the brain-extracted T1w using fast [FSL 5.0.9, RRID:SCR_002823, 84]. A T1w-reference map was computed after registration of 2 T1w images (after INU-correction) using mri_robust_template [FreeSurfer 6.0.1, 85]. Brain surfaces were reconstructed using recon-all [FreeSurfer 6.0.1, RRID:SCR_001847, 86], and the brain mask estimated previously was refined with a custom variation of the method to reconcile ANTs-derived and FreeSurfer-derived segmentations of the cortical gray-matter of Mindboggle [RRID:SCR_002438, 87]. Volume-based spatial normalization to two standard spaces (MNI152NLin6Asym, MNI152NLin2009cAsym) was performed through nonlinear registration with antsRegistration (ANTs 2.2.0), using brain-extracted versions of both T1w reference and the T1w template. The following templates were selected for spatial normalization: *FSL’s MNI ICBM 152 nonlinear 6th Generation Asymmetric Average Brain Stereotaxic Registration Model* [[88], RRID:SCR_002823; TemplateFlow ID: MNI152NLin6Asym], *ICBM 152 Nonlinear Asymmetrical template version 2009c* [[89], RRID:SCR_008796; TemplateFlow ID: MNI152NLin2009cAsym].

#### Functional data preprocessing

For each of the BOLD runs per subject (across all tasks and sessions), the following preprocessing was performed. First, a reference volume and its skull-stripped version were generated using a custom methodology of *fMRIPrep*. The BOLD reference was then co-registered to the T1w reference using bbregister (FreeSurfer) which implements boundary-based registration [90]. Co-registration was configured with nine degrees of freedom to account for distortions remaining in the BOLD reference. Head-motion parameters with respect to the BOLD reference (transformation matrices, and six corresponding rotation and translation parameters) are estimated before any spatiotemporal filtering using mcflirt [FSL 5.0.9, 91]. BOLD runs were slice-time corrected using 3dTshift from AFNI 20160207 [92, RRID:SCR_005927]. The BOLD time-series, were resampled to surfaces on the following spaces: *fsaverage*. The BOLD time-series (including slice-timing correction when applied) were resampled onto their original, native space by applying a single, composite transform to correct for head-motion and susceptibility distortions. These resampled BOLD time-series will be referred to as *preprocessed BOLD in original space*, or just *preprocessed BOLD*. The BOLD time-series were resampled into several standard spaces, correspondingly generating the following *spatially-normalized, preprocessed BOLD runs*: MNI152NLin6Asym, MNI152NLin2009cAsym. First, a reference volume and its skull-stripped version were generated using a custom methodology of *fMRIPrep*. Automatic removal of motion artifacts using independent component analysis [ICA-AROMA, 93] was performed on the *preprocessed BOLD on MNI space* time-series after removal of non-steady state volumes and spatial smoothing with an isotropic, Gaussian kernel of 6mm FWHM (full-width half-maximum). Corresponding “non-aggresively” denoised runs were produced after such smoothing. Additionally, the “aggressive” noise-regressors were collected and placed in the corresponding confounds file. Several confounding time-series were calculated based on the *preprocessed BOLD*: framewise displacement (FD), DVARS and three region-wise global signals. FD and DVARS are calculated for each functional run, both using their implementations in *Nipype* [following the definitions by 94]. The three global signals are extracted within the CSF, the WM, and the whole-brain masks. Additionally, a set of physiological regressors were extracted to allow for component-based noise correction [*CompCor*, 95]. Principal components are estimated after high-pass filtering the *preprocessed BOLD* time-series (using a discrete cosine filter with 128s cut-off) for the two *CompCor* variants: temporal (tCompCor) and anatomical (aCompCor). tCompCor components are then calculated from the top 5% variable voxels within a mask covering the subcortical regions. This subcortical mask is obtained by heavily eroding the brain mask, which ensures it does not include cortical GM regions. For aCompCor, components are calculated within the intersection of the aforementioned mask and the union of CSF and WM masks calculated in T1w space, after their projection to the native space of each functional run (using the inverse BOLD-to-T1w transformation). Components are also calculated separately within the WM and CSF masks. For each CompCor decomposition, the *k* components with the largest singular values are retained, such that the retained components’ time series are sufficient to explain 50 percent of variance across the nuisance mask (CSF, WM, combined, or temporal). The remaining components are dropped from consideration. The head-motion estimates calculated in the correction step were also placed within the corresponding confounds file. The confound time series derived from head motion estimates and global signals were expanded with the inclusion of temporal derivatives and quadratic terms for each [96]. Frames that exceeded a threshold of 0.5 mm FD or 1.5 standardised DVARS were annotated as motion outliers. All resamplings can be performed with *a single interpolation step* by composing all the pertinent transformations (i.e. head-motion transform matrices, susceptibility distortion correction when available, and co-registrations to anatomical and output spaces). Gridded (volumetric) resamplings were performed using antsApplyTransforms (ANTs), configured with Lanczos interpolation to minimize the smoothing effects of other kernels [97]. Non-gridded (surface) resamplings were performed using mri_vol2surf (FreeSurfer).

Many internal operations of *fMRIPrep* use *Nilearn* 0.5.2 [98, RRID:SCR_001362], mostly within the functional processing workflow. For more details of the pipeline, see the section corresponding to workflows in *fMRIPrep*’s documentation.

#### 3.8.1 ROI extraction

Regions of interest were identified using separate parcellations for the cerebellum, basal ganglia, and cortex. For the cerebellum, we extracted all regions using the atlas of Diedrichsen [25]; For the basal ganglia, we extracted the caudate, putamen, accumbens, and pallidum, using the Harvard Oxford atlas [99, 100]. Finally, for cortex, we extracted regions using the 400 region Schaefer parcellation [23]. For the Somatomotor A and B networks, we extracted only those regions which [23] identified as being activated during finger and tongue movement. For all regions and networks, we created timeseries data by computing the mean BOLD signal within each ROI.

#### 3.8.2 Covariance estimation and centering

For each subject, we used the Ledoit and Wolf shrinkage estimator [101] to derive covariance matrices for periods of equivalent length (100 imaging volumes) during baseline, and the beginning and end of the learning blocks of the VMR. For the the baseline blocks, we extracted 100 imaging volumes equally spaced throughout the scan, so as the capture the entirety of the baseline epoch of the task. For the reward-based motor task, we derived covariance matrices for the pre-learning, learning and post-learning epochs based on….

To center the covariance matrices, we took the approach advocated by [102], which leverages the natural geometry of the space of covariance matrices. We have implemented many of the computations required to replicate the analysis in an publicly available R package **spdm**, which is freely available from a Git repository at https://github.com/areshenk-rpackages/spdm.

The procedure is as follows. Letting *S_ij_* denote the *j*’th covariance matrix for subject *i*, we computed the grand mean covariance *S̅* over all subjects using the fixed-point algorithm described by [103], as well as subject means *S̅_i_*. We then projected the each covariance matrix *S_ij_*onto the tangent space at *S̅_i_* to obtain a tangent vector

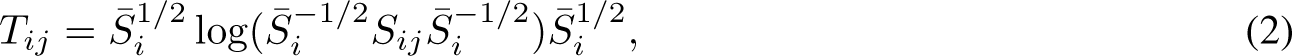

where log denotes the matrix logarithm. The tangent vector *T_ij_*then encodes the *difference* in covariance between the covariance *S_ij_* and the subject mean *S̅_i_*. We then transported each tangent vector to the grand mean *S̅* using the transport proposed by [102], obtaining a centered tangent vector *T^c^* given by

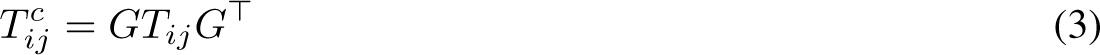

where 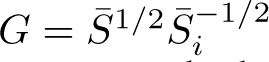. This centered tangent vector (a symmetric matrix) encodes the same difference in covariance, but now expressed relative to the mean resting state scan.

#### 3.8.3 Construction of the VMR learning-related neural axes and prediction of reward-based motor performance

Here we describe our approach for the main analysis – i.e., construction of the (performance corrected) explicit neural axis derived from the contralateral hand area, during the early learning epoch.

For each tangent vector in the early learning epoch, we computed the mean connectivity between the contralateral hand area (within the somatomotor network) and the remaining ROIs. The resulting vectors (containing an entry for each non-motor ROI) were used as the features for our predictive model. Call the resulting matrix of predictors **X** = [**x_1_**, **x_2_***, . . . ,* **x_n_**], where **x_i_** is a vector containing the mean connectivity of the *i*’th ROI with the hand area of motor cortex. We then computed the explicit learning score for each subject, defined as the circular mean difference between the target location and the subject’s reported aim direction. These scores formed the response variable **y**. Our *corrected* scores were then derived by orthogonalizing with respect to subjects’ VMR performance scores. That is, we regressed **y** onto the vector of VMR task scores and used the residuals for our analysis, and the same was done for each predictor (that is, each column in **X**). Both the predictors and outcome were then standardized prior to analysis.

We then fit a ridge regression model with separate network-level penalty terms using the model described by [28], and implemented in the R package GRridge [104]. That is, each of the 17 networks comprising the predictors – 15 non-motor cortical networks, plus cerebellum and basal ganglia – possessed their own shrinkage parameters. These parameters were separately tuned using an internal leave-one-out cross-validation loop, so that networks whose regions contributed relatively little to overall predictive performance were shrunken more harshly relative to predictively useful networks. The precise arguments were as follows:

grridge(X, y, partitions = network_labels, unpenal = ∼1, offset = NULL, method = “stable”, niter = 10,

innfold = NULL, fixedfoldsinn = TRUE, selectionEN = F, cvlmarg = 1, savepredobj = “all”)

The resulting parameter vector ***β*** contains, for each ROI, its estimated conditional contribution to subjects’ explicit report.

We then derived, for each subject, the mean functional connectivity of each region with the motor cortex during the learning period of the reward-based motor task (derived from the centered tangent vectors, as described above). The Pearson correlation between this vector and the vector ***β*** was the subject’s *explicit neural axis score*.

Note that all other models (such as the derivation of the implicit or performance neural axis), or neural axes using separate areas of motor cortex (e.g., tongue areas) or different time windows (e.g., baseline or pre-learning) were constructed identically, *mutatis mutandis* (e.g. using implicit scores rather than explicit reports).

